# IMPALA: A Comprehensive Pipeline for Detecting and Elucidating Mechanisms of Allele Specific Expression in Cancer

**DOI:** 10.1101/2023.09.11.555771

**Authors:** Glenn Chang, Vanessa L. Porter, Kieran O’Neill, Luka Culibrk, Vahid Akbari, Marco A. Marra, Steven J. M. Jones

## Abstract

**Summary:** Allele-specific expression (ASE), where transcripts from one allele are more abundant than transcripts from the other, can arise from various genetic mechanisms and has implications for gene regulation and disease. We present IMPALA (Integrated Mapping and Profiling of Allelically-expressed Loci with Annotations), a versioned and containerized pipeline for detecting ASE in samples including cancer genomes. IMPALA leverages RNA sequencing data and, optionally, phased variant, copy number variant (CNV), allelic methylation, and mutation data to identify ASE genes and uncover underlying regulatory mechanisms. IMPALA incorporates the MBASED framework for ASE detection, and outputs a comprehensive summary table and informative figures to visualize the genomic distribution of ASE genes and their correlation with potential regulatory causes. We applied IMPALA to a cancer sample and identified thousands of genes with ASE and highlighted potential somatic events that may have influenced ASE of these genes. ASE data can be used to detect the downstream consequences of genomic alterations, which facilitates the identification of dysregulated cancer-related genes. IMPALA thus provides researchers with a powerful tool for both ASE analysis and for investigating genetic factors correlated with ASE.

**Availability and implementation:** IMPALA is licensed under GNU General Public License v3.0 and freely available at https://github.com/bcgsc/IMPALA and https://doi.org/10.5281/zenodo.8019168 with documentation and tutorial.

**Supplemental information:** Supplemental materials are available at Bioinformatics online. Issue section: Gene expression

## Introduction

In diploid genomes, the majority of gene expression levels are balanced between the two alleles (Knight 2004; Cleary and Seoighe 2021). Allele-specific expression (ASE) is the imbalance in gene transcript abundance between two haplotypes (Knight 2004). ASE can be caused by various regulatory mechanisms that target single alleles (Robles-Espinoza *et al*. 2021). Examples of such mechanisms include epigenetic imprinting, allelically imbalanced copy number variants, and nonsense mutation on one allele that leads to nonsense-mediated decay of the transcript (Przytycki and Singh 2020; Robles-Espinoza *et al*. 2021). In cancer genomes, which are often characterized by high levels of genomic instability and epigenomic changes, dysregulation of ASE has been shown to play a role in the aberrant expression of cancer driver genes (Przytycki and Singh 2020).

To identify ASE, heterozygous single nucleotide variants (SNVs) within transcribed regions can distinguish the two alleles (Romanel 2019). ASE can be detected by examining allelic read counts at these SNVs in RNA sequencing (RNA-seq) data (Romanel 2019; Robles-Espinoza *et al*. 2021). A consensus can be created using multiple SNVs within a gene to obtain gene-level ASE data (Mayba *et al*. 2014).

Accurate genome phasing is beneficial for combining the allelic counts from multiple SNVs across a gene (Mayba *et al*. 2014; Wang *et al*. 2020). Long-read sequencing generates reads spanning over 10,000 base pairs (Martin *et al*. 2016; Wang *et al*. 2020), allowing multiple SNVs to be present within a single read, therefore generating longer phase blocks and more accurate haplotype phasing (Martin *et al*. 2016).

We present IMPALA (Integrated Mapping and Profiling of Allelically-expressed Loci with Annotations), a pipeline for detecting ASE in cancer. IMPALA takes a RNA alignment BAM file, an expression matrix, and an optional phased VCF file, and then generates a comprehensive table of major expressing allele frequencies for each expressed gene and the adjusted p-value of the gene displaying ASE (Fig 1A). The IMPALA pipeline also uniquely accepts optional genomic files to elucidate the underlying mechanisms of ASE. These data types include copy number variant data (Culibrk *et al*. 2021), allelic methylation data (Akbari *et al*. 2021), and a somatic mutation VCF as optional inputs. The IMPALA pipeline is fully containerized in a Snakemake workflow (Köster and Rahmann 2018) and can be executed using just a single command with appropriate modifications to the configuration files. The resulting ASE data can be used to confirm downstream effects of various genetic mechanisms and provide transcriptomic evidence of genomic alteration of cancer genes.

**Fig. 1.**
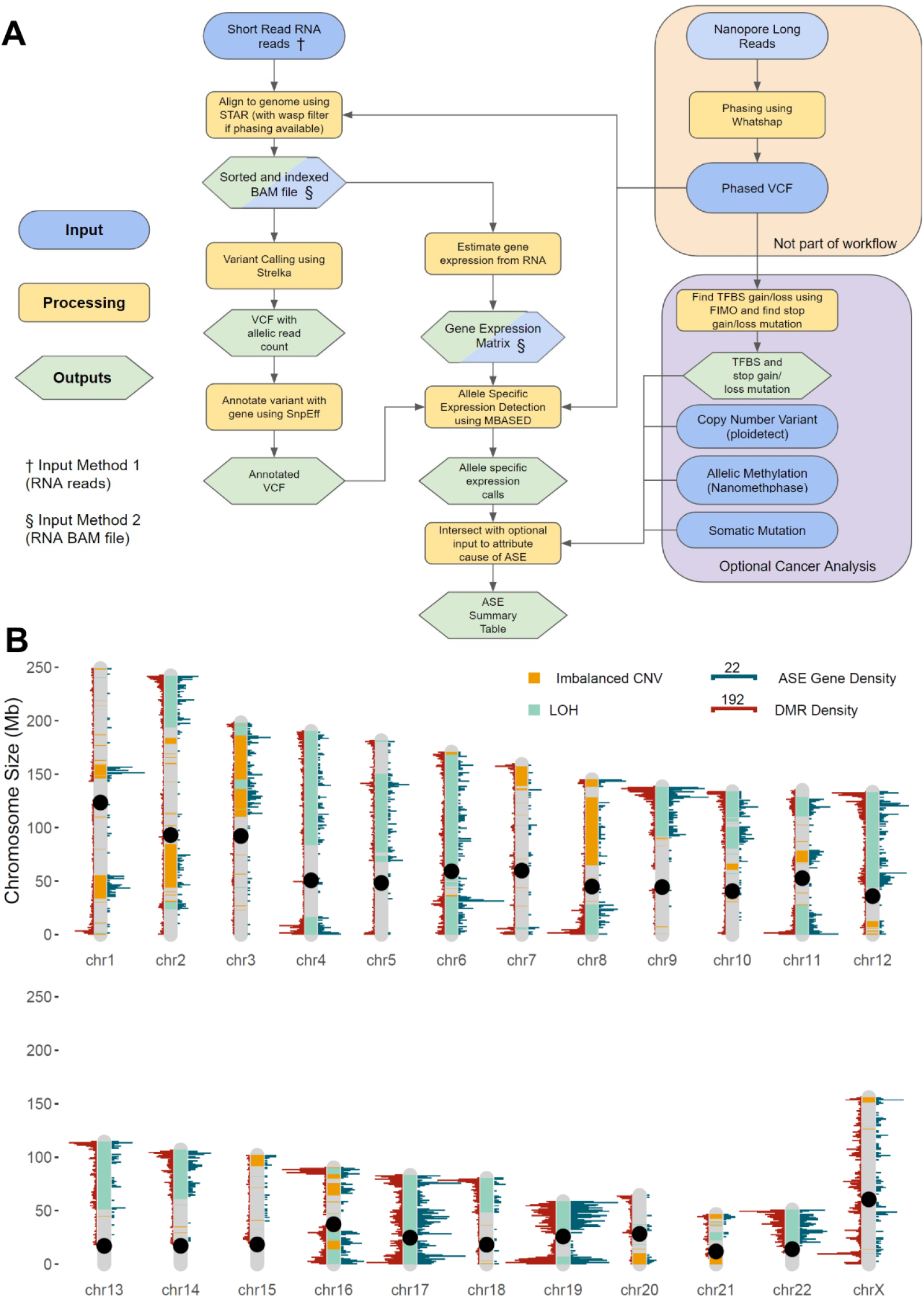
Overview of IMPALA. (A) General workflow showing inputs, processing steps and outputs of IMPALA. (B) Example karyogram figure automatically generated by IMPALA showing region with ASE genes (blue histogram on right), regions with DMR (red histogram on left) and allelic copy number variant state (color of chromosome band). The scales for the density of ASE genes and DMRs per 1 megabase region are indicated at the top and the black circle represents the centromeres.

## Methodology

### Genomic Phasing

We highly recommend using a phased VCF as input to IMPALA if long-read whole genome sequencing or trio data is available. The phased VCF file assigns SNVs within a haplotype block to an allele. Pseudo-phasing can be used as a substitution for inferring haplotypes but may yield a higher false positive rate. When analyzing ASE in cancer genomes, read-backed phasing is ideal as it leverages long-read data to phase the genome, including somatic variants above a detectable variant allele frequency (Roberts *et al*. 2021). We used Clair3 (v.0.1-r8) with the WhatsHap (v.1.0) phasing option for variant calling and phasing (Martin *et al*. 2016; Zheng *et al*. 2022).

### IMPALA preprocessing steps

The IMPALA workflow begins with aligning short paired-end RNA reads to a reference genome (Dobin *et al*. 2013). To minimize allelic mapping biases, the STAR (v. 2.7.10a) aligner employs the WASP parameter to filter out reads that do not map equally to the reference and non-reference genome (Van De Geijn *et al*. 2015). Variant calling on the resulting RNA BAM file is performed using Strelka (v. 2.9.10), enabling the extraction of allelic read counts (Kim *et al*. 2018). Only heterozygous SNVs are used, and if a phased VCF is provided, only heterozygous SNVs with phasing information are used. The genes affiliated with the filtered SNVs are then annotated using SnpEff (v. 5.0) (Cingolani *et al*. 2012b) and formatted with SnpSift (v. 5.1d) (Cingolani *et al*. 2012a). An expression matrix is also required for IMPALA to filter out genes with low expression. We generated our expression matrix from the RNA alignments using RSEM (v. 1.3.3) (Li and Dewey 2011).

### ASE calling (MBASED)

IMPALA uses the MBASED (v. 1.34.0) framework to quantify gene-level ASE using the annotated allelic read count, and if available, the phased VCF (Mayba *et al*. 2014). MBASED detects ASE in genes by conducting a beta-binomial test for each SNV and aggregating SNV level data to gene-level data using meta-analysis statistical framework and haplotype data (Mayba *et al*. 2014). When phasing information is available, SNVs are grouped by their specified haplotype. In the absence of phasing information, MBASED employs a pseudo-phasing algorithm, which assigns the allele with a higher read count at each SNV to one haplotype and the allele with a lower read count to the other haplotype (Mayba *et al*. 2014). The pseudo-phasing algorithm assumes that the direction of ASE is consistent among every SNV in a gene, which is not always true (Mayba *et al*. 2014).

### Cancer analysis - CNV

IMPALA can incorporate CNV data estimated from aligned tumor and normal DNA samples using the Ploidetect (v. 1.4.1) software (Culibrk *et al*. 2021). The CNV data allows IMPALA to categorize chromosomal regions into three CNV states: allelically balanced, allelically imbalanced, or regions with loss of heterozygosity (LOH). In balanced regions, both alleles have the same copy number, while imbalanced regions have different copy numbers on each allele. LOH regions indicate the complete loss of one allele. The CNV data are intersected with ASE results from MBASED using bedtools (v. 2.23.0) (Quinlan and Hall 2010). When CNV information is provided, the final output summary table includes four additional columns with the copy number for both alleles, the CNV state, and the expected major allele frequency for each gene based on the copy number.

### Cancer analysis - Allelic Methylation

The NanoMethPhase software (v. 1.2.0) uses Oxford Nanopore Technology long reads to obtain allelic methylation information, which can be used as input for IMPALA (Akbari *et al*. 2021).

Alternatively, methylation data can be directly obtained from PacBio long reads which can subsequently be phased and undergo differential methylation analysis to obtain allelic methylation calls. Using the allelic methylation calls, IMPALA searches for differential methylation between the alleles in the promoter region of each gene, spanning 2000 base pairs upstream and 500 base pairs downstream of the gene. It then determines instances where allelic methylation may be responsible for silencing one allele. It is essential to use the same phasing data for both IMPALA and NanoMethPhase to ensure consistent haplotype calling in the methylation data and the ASE data. With the addition of this parameter, the difference in allelic methylation at the gene promoter is added as an additional column within the summary table.

### Cancer analysis - Nonsense mediated decay

IMPALA can also optionally incorporate somatic SNV and Indel (Insertion/Deletion) information. The Samtools (v. 1.16.1) intersect tool is employed to identify somatic mutations within genes and their promoter regions (Danecek *et al*. 2021). The presence or absence of somatic mutations within genes and their promoters are then relayed in a Boolean column within the summary table.

When a phased VCF is provided, IMPALA also identifies nonsense mutations as potential correlates of ASE by the mechanism of NMD of the affected transcript. Heterozygous nonsense mutations are searched for by annotating the phased VCF using snpEff (v. 5.0) to determine the variant’s effect (Cingolani *et al*. 2012b). The nonsense mutation is expected to occur on the minor expressing allele as those transcripts decay via NMD. A column is added to the summary table whether a gene contains a heterozygous nonsense mutation.

### Cancer analysis - Transcription factor binding site mutation

IMPALA can also search for transcription factor binding site mutations in the promoter region of genes using the phased VCF and the reference genome fasta sequence. A consensus sequence for each allele is generated by incorporating phased variants into the reference genome using bcftools’ consensus command (Danecek *et al*. 2021). The FIMO software from the MEME suite (Grant, Bailey and Noble 2011) can then be used to scan for transcription factor motifs in the promoter region of each gene for both alleles. IMPALA filters out motifs with low expression (<1 TPM) of transcription factors and then identifies transcription factor motifs that differ between alleles. The information on transcription factor motifs is added to the summary table. It is important to note that experimental data are necessary to confirm whether a transcription factor is actually binding the gene promoter since FIMO uses computational methods to identify these sites.

### IMPALA output

The main output of IMPALA is a table that summarizes the key features of each gene, including its expression level, major allele frequency, copy number variant state, allelic methylation state, mutations, including presence of transcription factor binding site mutations, if all optional inputs are provided. As a point of reference, the table also includes the major allele frequency of each gene found in normal tissue, derived from the GTEx ASE data (Castel *et al*. 2020). IMPALA also automatically generates figures that visualize the distribution of the major allele frequency (Supplementary Fig. S1) as well as correlation with various genetic mechanisms (Fig 1B, Supplementary Fig. S2) to identify the mechanisms of ASE (Wickham *et al*. 2019).

#### Case Study (EGAD00001005859)

By combining evidence with various genetic mechanisms, we were able to use IMPALA to identify *CREBBP, INPP5D, NSD2, IRF1, TERT, ATRX*, and *STAG2* as candidate aberrant ASE genes involved in driving cancer in a tumor sample obtained from ovarian tissue of a patient diagnosed with ovarian adenocarcinoma. The genomic and transcriptomic data of this tumor sample can be downloaded from this EGAD00001005859 dataset. IMPALA identified 3,771 genes with significant ASE in the tumor sample (Supplemental Figure S1A). Notably, in this female patient sample, 157/184 (85.3%) of phased chromosome X genes exhibited ASE, presumably due to X-inactivation.

Our analysis revealed that allelically imbalanced CNV is the primary mechanism driving ASE in this sample, particularly in regions with LOH. Specifically, 2,491/3,771 (66.1%) of ASE genes were located in LOH regions compared to only 62/1,467 (4.2%) significant biallelic expressed (BAE) genes in this sample (Supplementary Fig. S2A). *CREBBP, INPP5D, NSD2* and *IRF1* are known cancer driver genes that showed LOH-mediated ASE. These genes also had somatic indel mutations on the major expressing allele, which may suggest a double-hit knockout mechanism. In contrast, copy number balanced regions exhibited a lower proportion of ASE, representing only 815/3,771 (21.6%) ASE genes, while 1,302/1,467 (88.8%) significant BAE genes were found to be CNV balanced (Supplementary Fig. S2A). Moreover, IMPALA identified 873/3,771 (23%) ASE genes with allelic methylation in the promoter, and among them, 775/873 genes (88.8%) exhibited allelic methylation silencing, where the allele with methylated promoter is in *trans* with the major expressing allele (Supplementary Fig. S2B). Notably, three of those genes are cancer-related, namely *TERT, ATRX*, and *STAG2*, where aberrant allelic methylation and expression may contribute to driving cancer. IMPALA also found 12 ASE genes that contain a nonsense mutation, 6 of which were on the minor expressing allele, which could be explained by nonsense mediated decay of transcripts originating from that allele (Supplementary Fig. S2C). This case study illustrates the use of IMPALA in identifying candidate ASE genes.

## Discussion

IMPALA offers a user-friendly workflow designed to efficiently identify and elucidate the potential mechanisms underlying ASE in a given sample. This pipeline is versioned and containerized, which greatly facilitates its distribution and reproducibility. Using an ovarian cancer sample as an example, genes with ASE have been demonstrated to exhibit correlations with allelic methylation and CNV, thus offering valuable insights into potential cancer driver genes within this sample. ASE data from IMPALA thus provides evidence of aberrantly-expressed genes to identify genes involved in driving cancer.

