## Supplemental Figures for "IMPALA: A Comprehensive Pipeline for Detecting and Elucidating Mechanisms of Allele Specific Expression in Cancer"

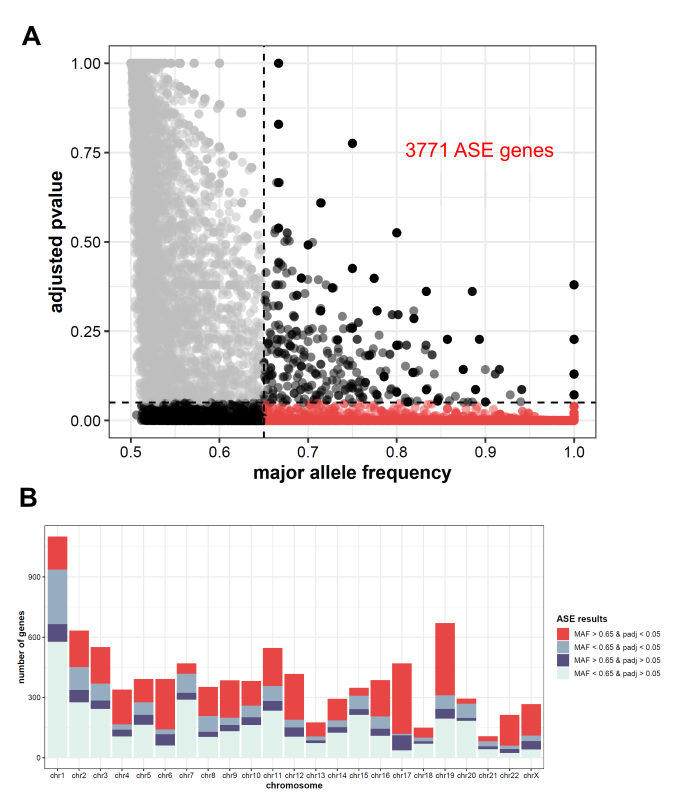

**Supplemental Figure S1.** Figures automatically generated from IMPALA after every run showing the distribution of ASE genes. **(A)** Scatterplot showing the major allele frequency and adjusted p-value (Benjamini-Hochberg procedure) of each gene calculated from MBASED. Genes with major allele frequency above a threshold (0.65 in this case) and p-value below 0.05 are considered ASE genes. 3,771 genes show ASE in this sample. **(B)** Bar graph showing frequency of genes along chromosomes. The color of the bar is classification of genes based on the major allele frequency threshold and p-value, where the red region is significant ASE genes.

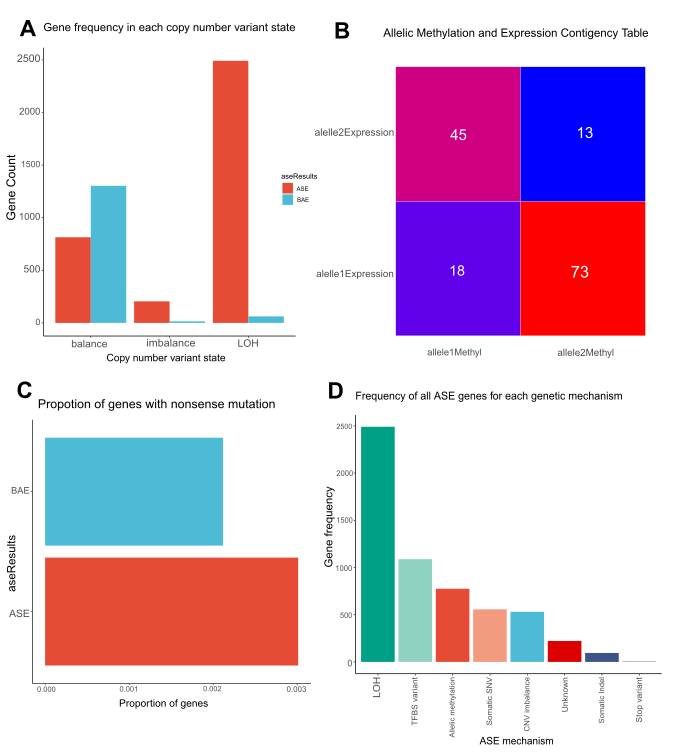

**Supplemental Figure S2.** Figures automatically generated from IMPALA based on optional genomic files to show correlation with cis-acting regulatory mechanisms. **(A)** Bar graph showing frequency of ASE and BAE genes in each allelic copy number variant state. Only generated when copy number variant data is provided. **(B)** Contingency graph showing ASE genes with allelic methylation. Genes are separated based on haplotype of allelic methylation and major expressing allele. Only generated when allelic methylation data is provided. **(C)** Bar graph showing proportion of genes in ASE and BAE that contain nonsense mutation. Only generated if phased VCF is provided. **(D)** Summary bar graph showing frequency of ASE that can be explained by each biological mechanism. The number of columns are based on genomic files provided. ASE genes that cannot be explained by any of the mechanisms are classified as unknown.
